# Adult and regenerating planarians respond differentially to chronic drug exposure

**DOI:** 10.1101/2022.08.02.502372

**Authors:** Kevin Bayingana, Danielle Ireland, Elizabeth Rosenthal, Christina Rabeler, Eva-Maria S. Collins

## Abstract

There is a lack of data on the effects of chronic exposure to common drugs and stimulants on the developing nervous system. Freshwater planarians have emerged as a useful invertebrate model amenable to high-throughput behavioral phenotyping to assay chemical safety in adult and developing brains. Here, we leverage the unique strength of the system to test in parallel for effects on the adult and developing nervous system, by screening ten common drugs and stimulants (forskolin, clenbuterol, LRE-1, MDL-12,330A, adenosine, caffeine, histamine, mianserin, fluoxetine and sertraline) using the asexual freshwater planarian *Dugesia japonica*. The compounds were tested up to 100 µM nominal concentration for their effects on planarian morphology and behavior. Quantitative phenotypic assessments were performed on days 7 and 12 of exposure using an automated screening platform. The antidepressants sertraline and fluoxetine were the most potent to induce lethality, with significant lethality observed at 10 µM. All ten compounds caused sublethal morphological and/or behavioral effects, with the most effects, in terms of potency and breadth of endpoints affected, seen with mianserin and fluoxetine. Four of the compounds (forskolin, clenbuterol, mianserin, and fluoxetine) were developmentally selective, causing effects at lower concentrations in regenerating planarians. Of these, fluoxetine showed the greatest differences between the two developmental stages, inducing many behavioral endpoints in regenerating planarians but only a few in adult planarians. While some of these behavioral effects may be due to neuroefficacy, these results substantiate the need for better evaluation of the safety of these common drugs on the developing nervous system.

## 1. Introduction

Over-the-counter (OTC) and prescription drugs are widely used to treat common ailments. According to data from the Centers for Disease Control and Prevention, about 46% of adults in the United States used prescription drugs in the last 30 days (2019). More than 90% of women take OTC drugs during pregnancy (Servey and Chang 2014) and most pregnant women use at least one prescription drug (Daw et al. 2011). Common OTC medications and prescription drugs used during pregnancy include antidepressants, antiemetics, antibiotics, analgesics, histamine receptor agonists and antagonists, heart medications, and cancer medication (Servey and Chang 2014; Haas et al. 2018). Antidepressant usage is more than twice as common in women than in men (CDC website). It is estimated that 2-3% of pregnant women take antidepressants during pregnancy (Dubovicky et al. 2017), with some longitudinal studies citing prevalence as high as 6% (Haas et al. 2018), raising concerns about possible side effects on the developing child. Selective serotonin reuptake inhibitors (SSRIs), such as fluoxetine and sertraline, are the most commonly used antidepressants during pregnancy (Cipriani et al. 2009; Dubovicky et al. 2017).

Antihistamine usage is also prevalent during pregnancy, to treat common allergies or dermatological symptoms like pruritus. About 10-15% of American women reported antihistamine use during pregnancy based on data from 1997 – 2011 (Hansen et al. 2020). Beta-adrenergic agonists are commonly used as bronchodilators as the treatment of choice for asthma, including during pregnancy (Billington et al. 2017). This drug class has also been used off-label to inhibit uterine contractions during pre-term labor, though the effectiveness of this treatment is unclear (The Canadian Preterm Labor Investigators Group 1992).

The safety of these drugs during pregnancy, however, remains unclear (So et al. 2010; Hansen et al. 2020) and usage of some drugs, such as the beta-adrenergic agonists, has been correlated with increased risk of neurological disorders, including Autism Spectrum Disorder (Witter et al. 2009). It is particularly difficult to ascertain the safety of these chemicals to the developing embryo because of the ethical concerns of human clinical trials with pregnant women. Thus, safety decisions are largely based on animal data that is expensive and time-consuming to obtain. Because of these limitations, the possible long-term effect that prenatal and early life exposure to these drugs has on the developing nervous system is largely understudied. Therefore, there is an urgent need for alternative high-throughput new approach methods to fill this data gap. High-throughput *in vitro* testing using mammalian or human cell lines has become a popular alternative for some toxicity assays, e.g., skin sensitization tests, and has successfully replaced traditional animal testing (OECD 2021). However, neurotoxicity and developmental neurotoxicity studies are especially difficult to assess *ex-vivo* (Bal-Price et al. 2010). Nervous system development and function depends on a complex network of signaling pathways spanning multiple cell types (e.g., neurons and glia), which is hard to recapitulate in 2-D culture systems. Therefore, non-mammalian organismal models have gained popularity for detecting systems-level adverse outcomes on the nervous system in a time- and cost-efficient manner (Peterson et al. 2008; Giacomotto and Ségalat 2010). Invertebrate models, such as fruit flies (Rand 2010; Chifiriuc et al. 2016), nematodes (Helmcke et al. 2010; Ruszkiewicz et al. 2018; Hunt et al. 2020), and freshwater flatworms (planarians) (Hagstrom et al. 2016; Wu and Li 2018), occupy a special role in this context. Due to their small size, they are amenable to high-throughput screening (HTS) in 48-, 96- or even 384-well plate formats, allowing for rapid screening of large chemical libraries (Ségalat 2007; Helmcke et al. 2010; Rand 2010; Giacomotto and Ségalat 2010; Hagstrom et al. 2016; Zhang et al. 2019a).

Fruit flies and nematodes, however, are difficult to dose with chemicals. Fruit flies cannot be grown in culture and nematodes have a cuticle which impedes chemical absorption (Giacomotto and Ségalat 2010; Kokel et al. 2012). In contrast, planarians are easily exposed to chemicals in their aquatic environment. Moreover, planarians have a wide array of robust phenotypic behaviors that are amenable to automated analysis and can be used for efficient functional screening of neurotoxicants (Zhang et al. 2019a; Ireland et al. 2020). Moreover, because of the similar size of adult and regenerating planarians, it is possible to screen adult and developing animals together using the same assays, enabling unprecedented studies comparing chemical toxicity in adults and during development (Hagstrom et al. 2015; Zhang et al. 2019a, b).

This unique strength allows us to evaluate whether chronic (12 day) exposure to commonly used drugs and stimulants differentially affects the developing versus the adult nervous system. We chose a set of ten compounds that are known to affect adenyl cyclase, beta-adrenergic receptor, adenosine receptor, histamine receptor, or 5-HT (serotonin) receptor activities in the mammalian brain (**Table 1**), pathways which are targeted by some common drugs as described above. The specific compounds were selected based on their possible use during pregnancy, including OTC (forskolin, caffeine) and prescription drugs (fluoxetine, sertraline, mianserin, clenbuterol, adenosine), or for their known use in research to target some of these pathways (LRE-1, MDL- 12,330A, histamine). Adenosine and caffeine were included because maternal cardiac arrhythmias are common during pregnancy and can be treated with intravenous injection of adenosine for acute termination of certain types of tachycardia (Ferrero et al. 2004; Joglar and Page 2012). Caffeine is an antagonist for the adenosine receptor and widely consumed among pregnant women in the U.S. (Chen et al. 2014). Moreover, several of the ten compounds have existing studies in flatworms (forskolin, caffeine, MDL-12, 330A, mianserin, histamine, fluoxetine, sertraline; **Supplemental Table**) enabling a direct comparison of our results with the literature.

**Table 1:** Chemical overview. The highest concentration of all chemicals tested was 100 µM. Water solubility information was obtained from the Sigma-Aldrich safety data sheet, DrugBank, or PubChem. Because activation of beta-adrenergic receptors subsequently upregulates adenyl cyclase, clenbuterol was grouped with the chemicals targeting adenyl cyclase. N/A: not available.

| Chemical | Mode of action | CAS # | Water solubility ($\mu\text{M}$) |
| --- | --- | --- | --- |
| Forskolin | Adenylyl cyclase agonist | 66575-29-9 | $2.7 \times 10^3$ |
| Clenbuterol hydrochloride | Beta-adrenergic receptor agonist | 21898-19-1 | $3.6 \times 10^2$ |
| LRE-1 | Adenylyl cyclase antagonist | 1252362-53-0 | N/A |
| MDL-12,330A hydrochloride | Adenylyl cyclase antagonist | 40297-09-4 | $8.0 \times 10^3$ |
| Adenosine | Adenosine receptor agonist | 58-61-7 | $5.2 \times 10^4$ |
| Caffeine | Adenosine receptor antagonist | 58-08-2 | $1.1 \times 10^5$ |
| Histamine dihydrochloride | Histamine receptor agonist | 56-92-8 | $9.0 \times 10^5$ |
| Mianserin hydrochloride | Histamine receptor antagonist | 21535-47-7 | $7.7 \times 10^2$ |
| Fluoxetine hydrochloride | Selective serotonin reuptake inhibitor | 56296-78-7 | $4.0 \times 10^4$ |
| Sertraline hydrochloride | Selective serotonin reuptake inhibitor | 79559-97-0 | 0.4 |
| L-ascorbic acid | Negative control | 50-81-7 | $1.4 \times 10^6$ |

Using an automated, robotic HTS system, we assayed a broad array of morphological and behavioral endpoints at two time points (days 7 and 12). We found that there were differential effects of some of these drugs, most notably mianserin and fluoxetine, both in terms of endpoints affected and in potency, in adult and regenerating planarians. Regenerating planarians were especially sensitive to fluoxetine exposure, warranting further studies into the vulnerable time for fluoxetine exposure and the molecular mechanisms by which fluoxetine may affect the developing brain.

## 2. Materials and methods

### 2.1. Planarian maintenance

We used asexual *D. japonica* planarians cultivated as described previously (Zhang et al. 2019a; Ireland et al. 2020). Planarians were reared in plastic containers with 0.5 g/L Instant Ocean (IO) water (Spectrum Brands, Blacksburg, VA, USA) and stored in the dark at 20°C when not used for experiments. Planarians were fed beef liver once or twice per week and cleaned the day of feeding and two days after feeding.

### 2.2. Chemicals

A list of the 10 drugs tested (including CAS number identifier and water solubility) is provided in **Table 1**. All chemicals were purchased from Sigma-Aldrich (St. Louis, MO, USA) and had a purity of ≥ 98%. Stock solutions were prepared in 100% dimethyl sulfoxide (DMSO, Sigma-Aldrich). Each drug was tested at a nominal concentration of 1, 3.16, 10, 31.6, and 100 µM, with a final DMSO concentration of 0.5%, which has no effect on the morphology or behavior of *D. japonica* (Hagstrom et al. 2015). Because significant effects were present in the lowest tested concentrations of fluoxetine and mianserin in regenerating planarians, these chemicals were rescreened in regenerating planarians at 0.00316, 0.01, 0.0316, 0.1, and 0.316 µM and at 0.316 µM, respectively. L-ascorbic acid (100 µM) was used as a negative control because it previously had no effect on any endpoint in our screening assay (Zhang et al. 2019a).

### 2.3. Chemical exposure

Screening plate set-up was performed as previously described (Zhang et al. 2019a). On the day of plate set-up (day 1), we randomly selected normally gliding worms that had been starved for at least 5 days. For adult planarians, we used worms of 5-10 mm length. For the regenerating planarians, slightly larger individuals were selected such that the final sizes of the amputated tail pieces were similar to that of the adult planarians. Regenerating worms were amputated below their auricles and above their pharynx with an ethanol-sterilized razor blade shortly before (< 3 hours) exposing them to the test compounds. The chemicals were tested on adult and regenerating worms separately, with the 5 concentrations and the 0.5% DMSO control in separate rows of each 48-well plate (Genesee Scientific, San Diego, CA; catalog # 25-108). The plates were sealed with ThermalSeal RTS seals (Excel Scientific, Victorville, CA) and kept in the dark, except for during screening. For each drug and concentration, we screened 3 replicates with n=8 planarians for each replicate, for a total n=24. For some chemicals, reruns were performed due to issues with vehicle control health or technical malfunctions, and thus n>24 for certain conditions. For each replicate, the orientation of the chemical concentrations was shifted to reduce the impact of edge effects (Zhang et al. 2019a).

### 2.4. Screening endpoints

Using a modified version of the fully automated HTS platform (Zhang et al. 2019a), we tested adult and regenerating worms using morphological and behavioral endpoints on day 7 and day 12 post-exposure. Our screening platform consists of a robotic microplate hander (Hudson Robotics, Springfield Township, NJ), and multiple cameras and assay stations as previously described (Zhang et al. 2019a).

Systemic toxicity endpoints consisted of lethality, body shape, eye regeneration (for regenerating worms only), and stickiness. Body shape was categorized both as the presence of any abnormal body shape and also separated out into distinct body shapes (e.g., lesion, contracted, C-shape, corkscrew, muscle waves) (Hagstrom et al. 2016; Ireland et al. 2020). Criteria used to distinguish the body shapes are provided in **Table 2** and example images are in **Figure 1**. Each planarian could be scored with up to 3 different abnormal body shapes. Behavioral endpoints consisted of measures of locomotion and spatial exploration, phototaxis, thermotaxis, and response to noxious heat, as described previously (Zhang et al. 2019a; Ireland et al. 2020, 2022). For all endpoints, data was analyzed using custom MATLAB scripts (MathWorks, Natick, MA). Any manual analyses were conducted by an investigator without knowledge of the identity of the drugs.

**Table 2:** Body shape definitions. Definitions included are only for shapes observed in this study.

| Body shape | Definition |
| --- | --- |
| Lesions | Loss of pigment, head regression or abnormal head shape, partial disintegration |
| Contracted | Shortened length, presence of ruffling along the exterior, immobile |
| C-shaped | C-shape, curled, or on its side |
| Corkscrew | Twisting around itself, screw-like hyperkinesia |
| Muscle Waves | Muscle waves, oscillation of length (e.g., peristalsis or scrunching (Cochet-Escartin et al. 2015)) |

**Figure 1.**
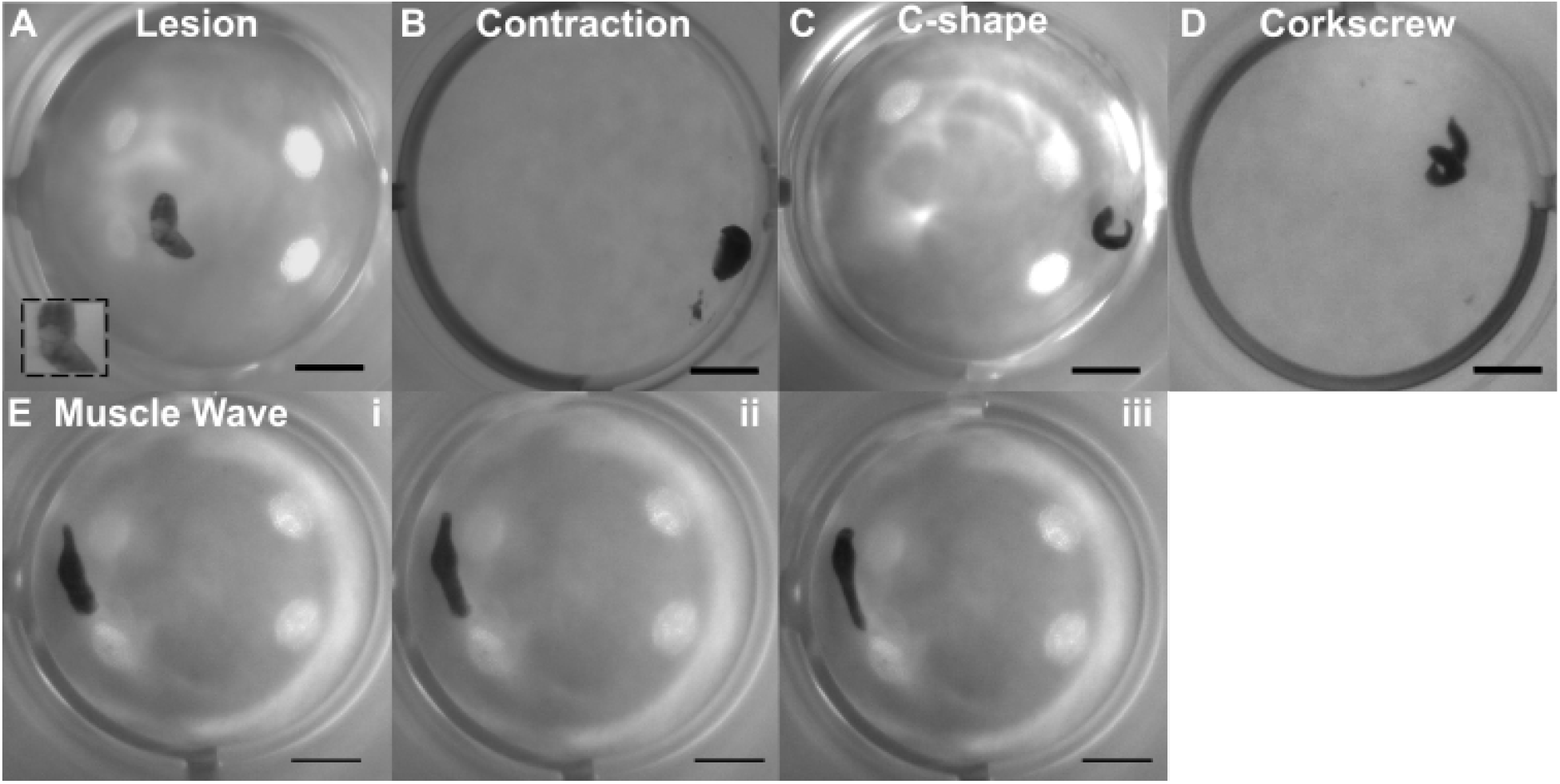
Examples of abnormal body shapes. (A-D) Example images of static body shapes: A) lesion, B) contracted, C) C-shape, and D) corkscrew. The inset in (A) shows a zoom-in of the lesioned area on the worm. E) Example image sequence of the dynamic muscle wave body shape. Muscle wave is shown as an oscillatory sequence of movement that begins from the head and propagates down the longitudinal axis of the planarian body. Scale bar: 2 mm. Brightness of the original images was adjusted as necessary to aid visualization. Exposure conditions: A: MDL-12, 330A; B, D, and E: mianserin; C: sertraline.

### 2.5. Statistical analysis

Statistical analysis was performed on compiled data from at least triplicate runs. Each concentration was compared with its 0.5% DMSO control as previously described (Zhang et al. 2019a). For lethality, eye regeneration, body shape, stickiness and scrunching endpoints, statistical significance was determined using one tailed Fisher’s exact test. For speed, resting, thermotaxis, phototaxis, noxious heat sensing, anxiety, and locomotor bursts, we used a parametric two-tailed t-test or a nonparametric two-tailed Mann-Whitney U test depending on whether the sample distribution was normal or not, respectively. Normality of the samples was determined using Lilliefor’s test. A p-value less than 0.05 was statistically significant. Speed endpoints consisted of the average speed in 30 second bins throughout the phototaxis assay. For the dark period, if a specific concentration of a drug caused a significant effect in at least half of the increments, then it was marked as a hit for the dark period. For the blue light period, a hit in either of the 30 second bins constituted a hit for the whole light period. This distinction between light and dark was made because we observed more variability in the control behavior during the dark period.

We excluded inconsistent hits where only one replicate was statistically significant while the other two were not, as in (Zhang et al. 2019a). **Figure 2** provides an overview of our statistical pipeline. We report the data using lowest observed effect levels (LOELs), indicative of the lowest concentration with a statistically significant hit.

**Figure 2:**
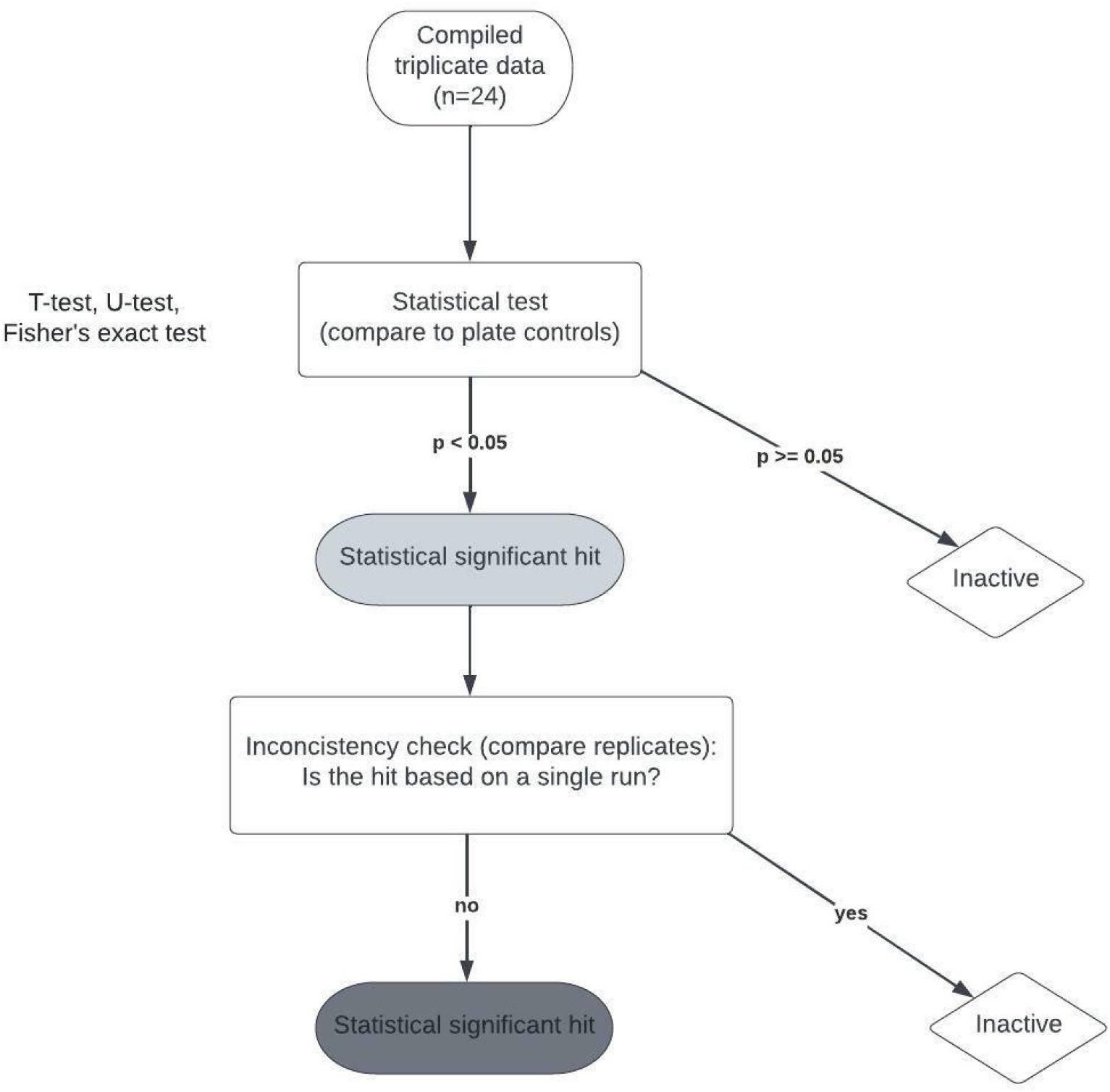
Workflow of statistical analysis.

### 2.6. Eye regeneration

To more closely evaluate effects on eye regeneration, regenerating planarians were exposed to mianserin and fluoxetine at 0.316, 1, 3.16 µM. The presence of eyes was manually scored every day until day 5. Duplicate experiments, each with 5 worms per condition, were performed for total of n=10 per condition.

### 2.7. Acetylcholinesterase activity

Thirty-six adult or regenerating planarians were exposed to either 0.5% DMSO solvent control, 3.16 µM fluoxetine, or 31.6 µM mianserin for 12 days. The planarians were maintained in 12-well plates (Genesee Scientific), with 6 planarians per well and a total volume of 1.2 mL of the test solution to keep the ratio of chemical/planarian consistent with the screening set-up. Any fission events or planarians from wells with death were excluded from the assay. Post-exposure, the planarians were washed 3X with IO water and homogenized in 1% Triton X-100 in PBS as described in (Hagstrom et al. 2017, 2018a). An Ellman assay (Ellman et al. 1961) was performed using an Acetylcholinesterase activity assay kit (Sigma-Aldrich). Absorbance was read at 412 nm every minute for 10 minutes using a VersaMax (Molecular Devices, San Jose, CA) spectrophotometer. Acetylcholinesterase activity was calculated as the rate of change of absorbance per minute during the linear portion of the reaction and normalized by protein concentration as determined by a Coomassie (Bradford) protein assay kit (Thermo Scientific, Waltham, MA). Activity was compared to the solvent control samples (set at 100% activity). Activity measurements were performed with at least three technical replicates and at least 2 independent experiments (biological replicates).

## 3. Results

### 3.1 Effects in adult planarians

Lethality was observed for all drugs in adult planarians in the tested concentration range (up to 100 µM), except for adenosine, caffeine, and histamine (**Figure 3A**). The tested SSRIs, fluoxetine and sertraline, showed the greatest toxicity, with significant lethality in adult planarians as low as 10 µM. Notably, sertraline has generally low water solubility (0.4 µM, **Table 1**), but as we observed lethality starting at 10 µM, a sufficient amount must be solubilized in our conditions, likely aided by the addition of DMSO. Mianserin was the only compound to cause abnormal body shapes (muscle waves and corkscrew) at day 7, which appeared with exposure as low as 1 µM. On day 12, abnormal body shapes (c-shapes) were also observed for adenosine at 100 µM. Increased stickiness was only observed in adult planarians exposed to 10 µM MDL-12,330A.

**Figure 3:**
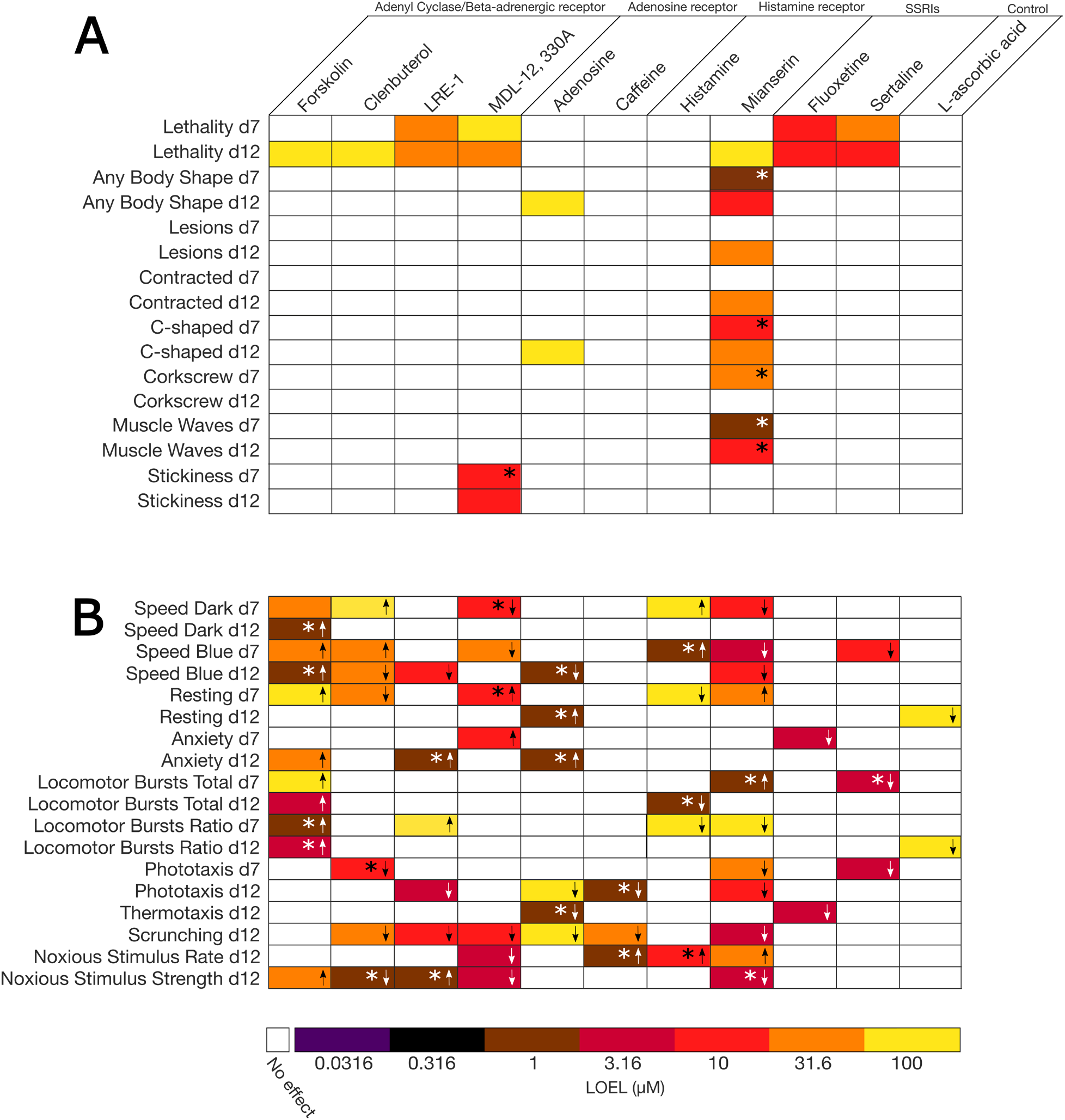
Heat map of effects in adult planarians. The lowest observed effect levels (LOELs) on days 7 (d7) and 12 (d12) of exposure are indicated for (A) systemic toxicity endpoints and (B) behavioral endpoints. For all endpoints besides lethality, only sublethal effects are shown. Drugs boxed are thought to act on the same receptor, with the drugs in the left-half of each box being agonists and the drugs in the right-half being receptor antagonists. The negative control L-ascorbic acid was only tested at 100 µM. * indicates concentration-independent effects. Arrows in (B) indicate the directionality of the effect compared to the vehicle controls.

All test compounds caused behavioral effects in adult planarians at sublethal concentrations (**Figure 3B**). Scrunching was one of the most sensitive endpoints which showed concentration-dependent hits. While hits were sometimes observed at lower concentrations in some of the other endpoints, they were often concentration-independent. When considering the locomotion-based endpoints (speed, resting), we observed both hypo- and hyper-active effects. Forskolin, clenbuterol, and histamine caused largely hyperactive effects (increased speed, decreased resting). Clenbuterol caused increased speed on day 7 but decreased speed on day 12, suggesting a potential change of mechanism over time. Other compounds, such as LRE-1, MDL-12,330A, adenosine, mianserin, and sertraline caused decreased speed and/or increased resting. Notably, the adenyl cyclase and beta-adrenergic receptor drugs showed opposing effects, with the agonists clenbuterol and forskolin causing hyperactivity, while the antagonists LRE-1 and MDL-12,330A caused hypoactivity. Additionally, these four drugs caused effects on scrunching and/or the strength of reaction to the heat (noxious stimuli strength), though the directionality of these effects was not correlated with the agonism/antagonism of the pathways.

The tested adenosine receptor drugs, adenosine and caffeine, caused relatively few hits, with significant effects only seen on day 12. Adenosine caused concentration-independent effects in speed in the blue period, anxiety and thermotaxis at 1 µM, as well as defects in phototaxis and scrunching at 100 µM. Similarly, with caffeine, concentration-dependent effects were only observed for scrunching at 31.6 µM. Mianserin caused hypoactive effects on many endpoints, with significant effects observed for 11/18 behavioral endpoints and effects on both day 7 and day 12, at concentrations as low as 1 µM for concentration-independent effects and 3.16 µM for concentration-dependent effects.

For the SSRIs, we observed relatively few behavioral effects and only at the highest sublethal concentration (3.16 µM). For fluoxetine, effects on both day 7 (anxiety) and day 12 (phototaxis) were observed, whereas sertraline caused effects only on day 7 (speed blue, total locomotor bursts and phototaxis).

### 3.2. Effects in regenerating planarians

Lethality was observed for forskolin, LRE-1, MDL-12,330A, mianserin, fluoxetine, and sertraline in regenerating planarians (**Figure 4A**). For forskolin, mianserin, and fluoxetine, lethality at day 12 was preceded by morphological effects (i.e., abnormal body shapes or eye regeneration defects) at day 7 at the same concentration. The lesions induced by forskolin at day 7 suggest that the planarians were getting sick before forskolin became significantly lethal at day 12. As in adult planarians, fluoxetine and sertraline showed the greatest toxicity, with significant lethality at 10 µM.

**Figure 4:**
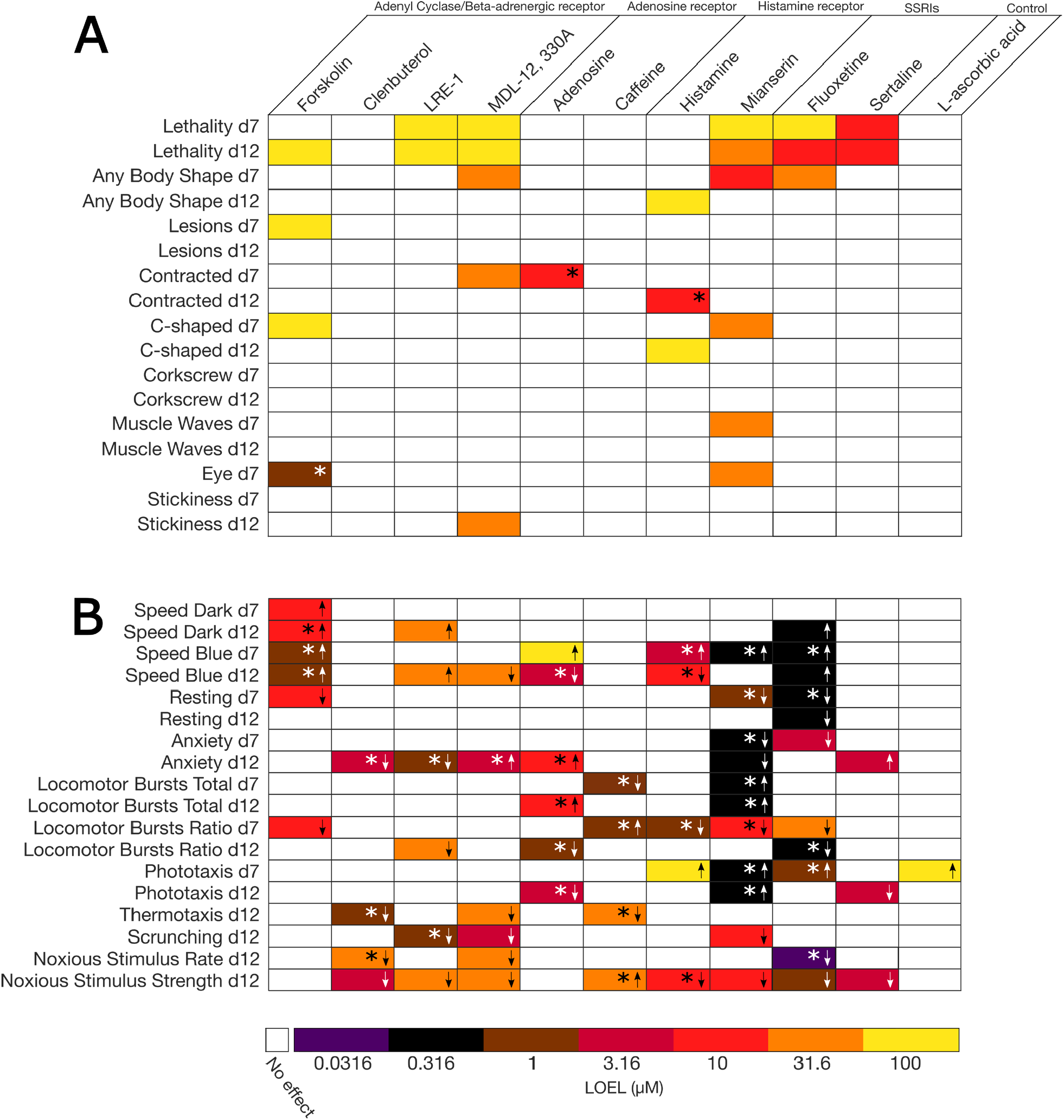
Heat map of effects in regenerating planarians. The lowest observed effect levels (LOELs) on days 7 (d7) and 12 (d12) of exposure are indicated for (A) systemic toxicity endpoints and (B) behavioral endpoints. For all endpoints besides lethality, only sublethal effects are shown. Drugs boxed are thought to act on the same receptor, with the drugs in the left-half of each box being agonists and the drugs in the right-half being receptor antagonists. The negative control L-ascorbic acid was only tested at 100 µM. * indicates concentration-independent effects. Arrows in (B) indicate the direction of effects in comparison to the vehicle controls.

Effects on planarian body shapes at sublethal concentrations were observed for MDL-12,330A, adenosine, histamine, and mianserin. Planarians exposed to 31.6 µM MDL-12,330A displayed contraction on day 7. Adenosine at 10 µM caused concentration-independent contracted body shapes at day 7. Mianserin induced abnormal body shapes as low as 1 µM (muscle waves), though this was concentration-independent. Concentration-dependent effects on shapes were observed starting at 10 µM mianserin and included c-shapes, corkscrew, and muscle waves. Histamine caused c-shapes at day 12 at 100 µM. MDL-12,330A caused increased stickiness on day 12.

All compounds caused sublethal behavioral effects in regenerating planarians (**Figure 4B**). Like in adult planarians, forskolin caused hyperactive effects on locomotion. However, unlike in adults, clenbuterol did not affect locomotion in regenerating planarians and LRE-1 caused hyperactive effects on day 12. Thus, the agonist/antagonist relationship seen with adult planarians was not found in the regenerating planarians. However, noxious heat sensation was still largely affected by the adenyl cyclase and beta-adrenergic receptor drugs as all compounds, except for forskolin, caused effects on scrunching and/or the noxious stimuli strength endpoint.

Adenosine and caffeine caused more hits in regenerating planarians than in adults, but these were mostly concentration-independent. Histamine similarly only caused concentration-independent effects or effects at the highest tested concentration (100 µM), whereas mianserin affected many behavioral endpoints, some of them at the lowest test concentrations. The most sensitive concentration-dependent effect was decreased anxiety on day 12, which was seen as low as 0.316 µM. All other effects at this concentration were concentration-independent, i.e., they did not show a monotonic concentration response. Fluoxetine similarly affected many behavioral endpoints with great sensitivity. Significant effects were seen as low as 0.0316 µM, though concentration-dependent hits were observed starting at 0.316 µM. Many of these sensitive hits represented hyperactive effects on locomotion. This is in stark contrast to what was observed in the adult planarians, where very few endpoints were affected and only starting at 3.16 µM. Unlike fluoxetine, sertraline caused similar effects in adults and regenerating planarians as few endpoints (anxiety, phototaxis, and noxious stimuli strength) were affected at the highest sublethal concentration (3.16 µM) and only at day 12. The breadth of effects, overall sensitivity, and differential sensitivity between developing and adult planarians seen with mianserin and fluoxetine prompted us to investigate these compounds in more detail.

### 3.3 Mianserin and fluoxetine

Mianserin greatly changed the behavior of adult and regenerating planarians and concentration appeared to play an important role in the manifestations of the different phenotypes. For example, a mix of different abnormal body shapes were observed in both adult and regenerating planarians and the predominant body shape was concentration-dependent (**Figure 5A**). In adult worms at day 7, low to mid concentrations of mianserin (1, 10 µM) induced primarily muscle waves. Starting at 31.6 µM, muscle waves were observed concomitantly with c-shapes and corkscrews. At 100 µM (which was lethal by day 12), the body shapes were predominantly c-shapes or a mix of c-shapes and muscle waves. On day 12, adult worms showed muscle waves at 10 µM and contraction and c-shapes at 31.6 µM. Similar patterns were seen with the regenerating planarians. On day 7, the occurrence of any body shape was statistically significant starting at 10 µM; however, no one body shape showed statistical significance as a low incidence of contraction, c-shapes, and muscle waves was observed. Like in adults, a mixture of c-shapes and muscle waves predominated at 31.6 µM (which was lethal at day 12). Unlike in the adult worms, only a low, non-statistically significant level of contraction was observed in the day 12 regenerating planarians.

**Figure 5:**
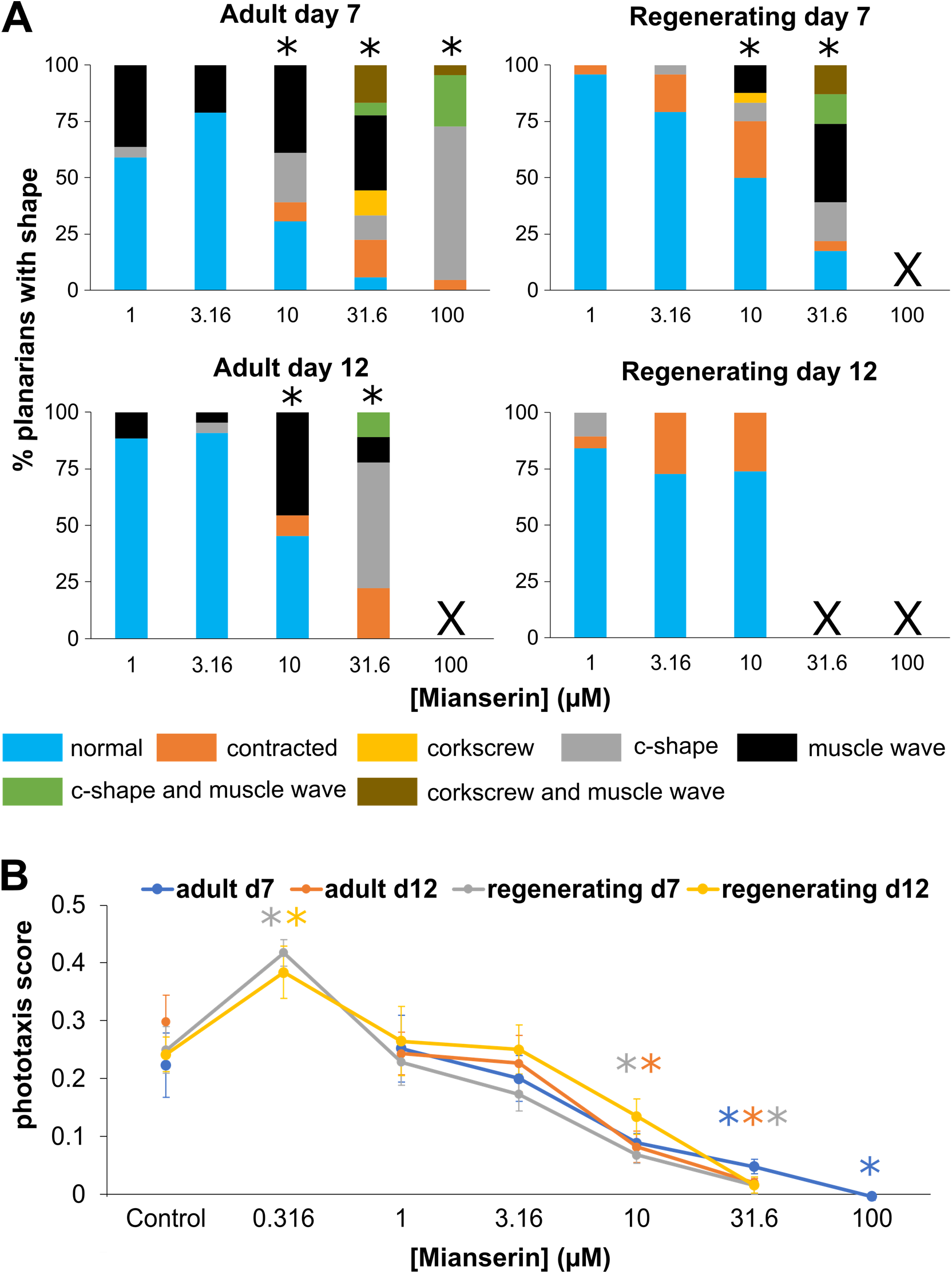
The effects of mianserin change with concentration. A) The percentage of abnormal body shapes observed at each concentration (µM) of mianserin is shown for adult and regenerating worms on days 7 and 12. No data are shown for concentrations with significant lethality (marked with X). B) Concentration-response curves of day 7 and day 12 phototaxis scores for adult and regenerating planarians exposed to mianserin. * indicates statistical significance (p<0.05).

Additionally, a non-monotonic concentration relationship was observed in day 7 phototaxis behavior for regenerating planarians exposed to mianserin (**Figure 5B**). At 0.316 µM, a statistically significant increase in phototaxis behavior was observed, whereas at 10 and 31.6 µM significant decreases in phototaxis behavior were observed. At day 12, the hyperactive effect at 0.316 µM was retained, but a decrease in phototaxis behavior was not seen at the higher concentrations. In adults, only hypoactive effects on phototaxis were observed. Similar trends were also observed for speed in the blue period.

Mianserin caused many effects at both developmental stages (**Figure 6A**), but the potency was different in the two worm types, with regenerating planarians displaying more sensitivity as they showed hits at lower concentrations compared to adult worms. As mentioned above, in regenerating planarians, mianserin exhibited both hyper- and hypo-active effects on locomotion, depending on the concentration, whereas only hypoactive locomotor effects were observed in the adults. In contrast, fluoxetine had minimal sublethal effects in adult worms - only lowering planarians’ anxiety and thermotaxis at the highest sublethal concentration (3.16 µM). However, fluoxetine affected many endpoints related to locomotion in regenerating worms (**Figure 6B**) and at much lower concentrations (as low as 0.0316 µM). Several of these effects were hyperactive, such as increased speed, decreased resting, and increased phototaxis. Although some of these effects were concentration-independent, some, such as increased speed in the dark and blue periods at day 12, were concentration-dependent with effects seen at 0.316, 1 and 3.16 µM.

**Figure 6:**
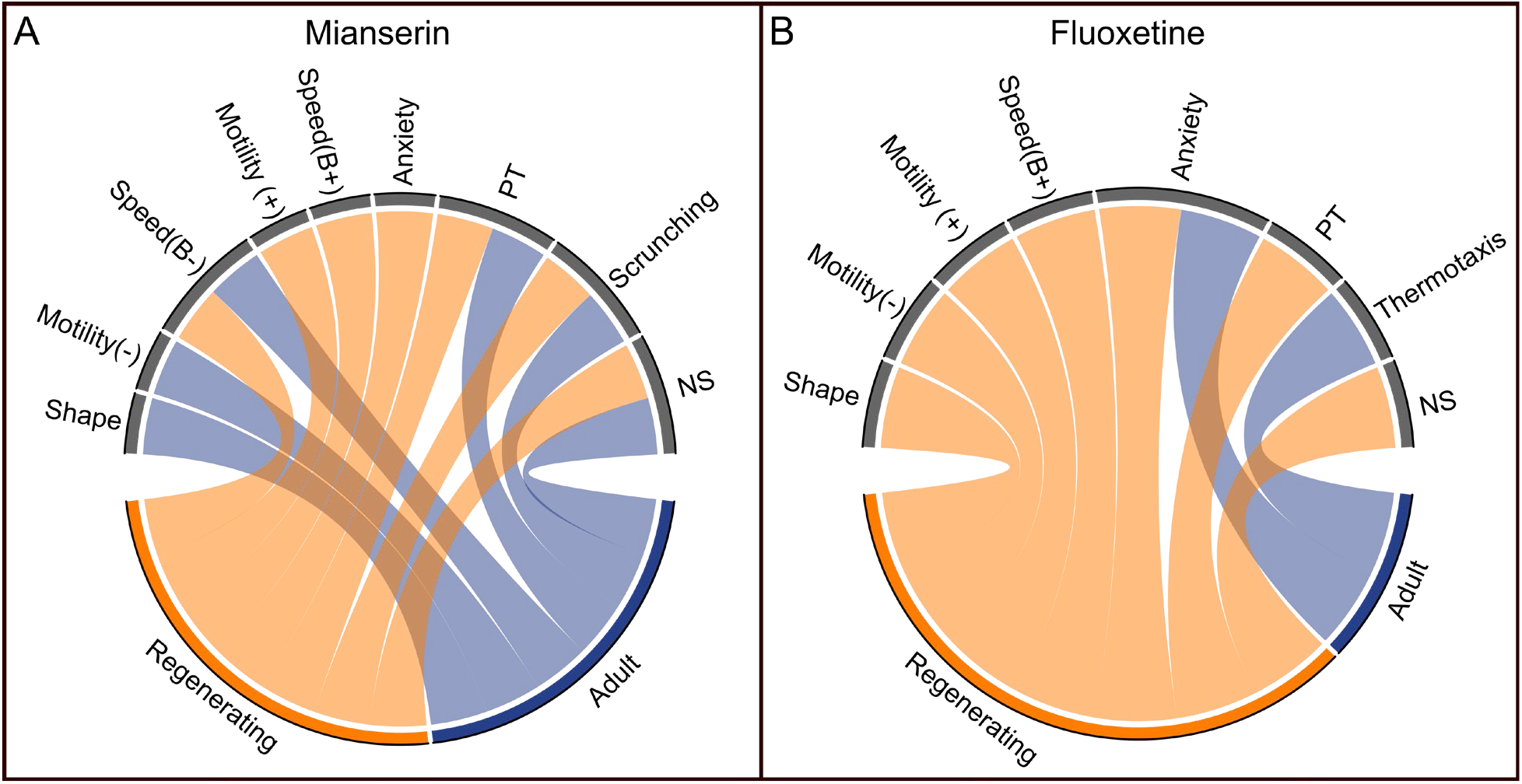
Differential effects of mianserin and fluoxetine on adult and regenerating planarians. Interaction of A) mianserin or B) fluoxetine with the different endpoint classes for adult and regenerating planarians. Connections were made if the chemical caused a hit at either day 7 or 12 at any tested concentration. Effects on speed in the dark period, resting, or locomotor bursts were combined into the “Motility” category. Speed(B): speed in the blue period, PT: Phototaxis, NS: noxious stimuli. For endpoints that had effects in either direction, either the positive (+) or negative (-) direction of the effects are noted.

*In vitro* studies have found that fluoxetine and other anti-depressants increase neuronal proliferation rates (Chang et al. 2009). Thus, because we cannot measure regeneration rates on our HTS platform, (though we can detect delayed eye regeneration at day 7, as observed with 31.6 µM mianserin), we assayed eye regeneration using stereo microscopy. Regenerating planarians were exposed to either mianserin or fluoxetine at 0.316, 1, or 3.16 µM, or to 0.5% DMSO (vehicle control) and eye regeneration was manually scored every day from days 1-5. All planarians had no eyes on day 4 and had 2 eyes on day 5, demonstrating no difference associated with chemical exposure (**Supplemental Figure 1A**).

Additionally, antidepressants have been found to alter acetylcholinesterase expression and/or function in a variety of animal systems (Yang et al. 2018), including humans (Müller et al. 2002). Therefore, we evaluated whether acetylcholinesterase activity was altered in adult or regenerating planarians exposed for 12 days to either 3.16 µM fluoxetine, or 31.6 µM mianserin. These concentrations were the highest sublethal tested concentrations in adult worms. No significant inhibition was observed at these concentrations (**Supplemental Figure 1B**).

## 4. Discussion and conclusions

It is well known that the developing brain is more sensitive to chemical exposure. However, we have little knowledge on the differential effects of compounds on the developing versus the adult brain, because studies in developing organisms are difficult. HTS using the planarian *D. japonica* is promising to fill this gap, as it allows for the parallel study of chemicals on adult and developing brains, using multiple quantitative morphological and behavioral endpoints. This allows us to detect specific developmental effects and distinguish them from general effects on the nervous system. Our screening work so far has focused on studying the effects of potential neurotoxicants on brain development and function; here, we study a group of 10 neuroactive drugs and stimulants, which we expected to show both desired neuroactive effects and neurotoxic effects. Caffeine, fluoxetine, and sertraline have known aquatic toxicity in the mM range (https://cfpub.epa.gov/ecotox/). However, reported toxicity in *Daphnia* is at least an order of magnitude higher than our highest test concentration (100 µM). Except for adenosine, caffeine, and histamine, we observed lethality at the highest 1-3 test concentrations for all compounds in regenerating and/or adult planarians, and any morphological or behavioral effects at these lethal concentrations are therefore likely due to systemic toxicity. Thus, we focused our analysis on the sublethal morphological and behavioral effects of these compounds.

### 4.1 Differential effects in adult versus regenerating planarians

All ten compounds produced behavioral changes at sublethal concentrations in both adult and regenerating planarians. Comparing outcomes for the two developmental stages, we discovered that many of the tested compounds caused differential effects in adult versus regenerating planarians. Lethality was generally induced at the same or lower concentrations in adult planarians than in regenerating planarians, similar to previously observed trends (Zhang et al. 2019a; Ireland et al. 2022). Only mianserin caused lethality at lower concentrations in regenerating planarians than in adults.

Looking at the sublethal effects on a per endpoint basis, we found that scrunching was a sensitive endpoint in adult planarians, with 6/10 compounds causing decreased scrunching in response to noxious heat. In contrast, only 3/10 compounds affected scrunching in regenerating planarians, with one hit being concentration-independent. Interestingly, regenerating planarians had more hits in the noxious stimulus strength measure, with 8/10 compounds showing effects (2 concentration-independent), whereas in adult planarians only 5/10 had effects, and three of those were concentration-independent. The interpretation of these differences is difficult because the molecular regulation of noxious heat sensation and the planarians’ behavioral responses are poorly understood. Cholinergic signaling seems to be involved in regulating planarian behavior in response to noxious heat because chemical and molecular inhibition of acetylcholinesterase can cause defects in these endpoints (Hagstrom et al. 2018b; Zhang et al. 2019b; Ireland et al. 2022). However, the identity of the heat sensitive receptors in *D. japonica* remains to be determined (Sabry et al. 2019) as well as how the initial noxious heat sensation is processed in the nervous system to cause the stereotypical muscle-driven periodic body length scrunching oscillations (Cochet-Escartin et al. 2015).

Regenerating planarians had increased sensitivity to four of the tested compounds (forskolin, clenbuterol, mianserin, and fluoxetine), showing lower overall LOELs than the adult planarians. The most striking example of this increased sensitivity was fluoxetine, which only showed decreased anxiety and thermotaxis in adult planarians at 3.16 µM, whereas it affected 12 non-lethality endpoints in regenerating planarians, with effects starting as low as 0.0316 µM (0.316 µM for concentration-dependent effects). The finding that both adult and regenerating planarians exposed to fluoxetine showed decreased anxiety may be related to fluoxetine’s action as an SSRI. The diversity in phenotypic profiles between the two developmental stages suggests that different targets may also be affected by fluoxetine in regenerating planarians or that the same targets have different roles during development versus in the adult organism. One such target may be acetylcholinesterase, which has been found to be inhibited by fluoxetine in other systems (Müller et al. 2002) and has been suggested to play a role in development, which may or may not rely on its enzymatic activity (Bigbee et al. 2000; Paraoanu et al. 2006; Layer et al. 2013). However, we did not observe significant inhibition of acetylcholinesterase at 31.6 µM fluoxetine, the highest sublethal tested concentration, in either adult or regenerating planarians. Notably, several of the effects in regenerating planarians were concentration-independent and/or indicative of hyperactivity, which may be the result of neuroactive and not neurotoxic effects of the drug. Since fluoxetine is approved for use in children as young as 8 years old to treat depression and bipolar disorder, the observed sensitivity of the regenerating worms suggests that further studies into possible adverse effects during neurodevelopment may be warranted.

### 4.2 HTS can recapitulate findings from low-throughput studies

Existing studies on these compounds in freshwater planarians or parasitic flatworms (*Schistosoma mansoni*) have been low-throughput and primarily based on manual scoring. Most studies focused on acute (≤ 3 hours) effects in adult worms, with a few exceptions that also investigated the effect of subacute (3 - 24 hours), short-term (2-3 days) or chronic (>3 days) exposure and/or on development and regeneration (reviewed in **Supplemental Table**). Inter- species comparisons with planarians can be challenging as different planarian species can have distinct sensitivities (Ireland et al. 2020) and even behavioral responses (Sabry et al. 2019) to the same chemical. However, comparisons are still useful to investigate whether HTS can recapitulate the general findings from low-throughput studies.

Decreases in motility have been found in adult flatworms acutely exposed to MDL- 12,330A (Matsuyama et al. 2004), mianserin (Currie and Pearson 2013; Talbot et al. 2014; Chan et al. 2016; Shettigar et al. 2021), sertraline (Thumé and Frizzo 2017; Weeks et al. 2018), and high concentrations of caffeine (Best and Morita 1982; Moustakas et al. 2015). Previous studies on the effects of caffeine in planarians have only found behavioral effects at much higher concentrations than we tested here (mM instead of µM). At these high concentrations, acute/subacute exposure to caffeine of different *Dugesia* species induced morphological defects (contractions and C-like hyperkinesia), paralysis, head lesions, and eventually death (Best and Morita 1982; Rawls et al. 2010; Li 2013; Moustakas et al. 2015). We found that chronic exposure to low concentrations caused few behavioral or morphological effects, suggesting that low to moderate concentrations are tolerated by adult and developing planarians. Notably, moderate consumption of up to 200 mg caffeine/day during pregnancy is considered safe by the American College of Obstetricians and Gynecologists (American College of Obstetricians 2010).

Acute exposure to mianserin has previously been shown to induce muscle-based movement in *Schmidtea mediterranea* planarians (Currie and Pearson 2013; Talbot et al. 2014). Our finding that low to mid-concentrations of mianserin cause a significant induction of muscle waves suggests that mianserin acts similarly in *D. japonica*. In addition to motility defects, exposure to 10 µM mianserin has been reported to induce regeneration of two-headed *D. japonica* planarians 6% of the time in amputated trunk pieces, suggesting weak effects on body axis polarity (Chan et al. 2014). Here, defects in eye regeneration at day 7 were observed with 31.6 µM, which causes significant lethality by day 12.

Acute to short-term exposure to sertraline has been reported to cause many morphological effects, including degeneration in the parasitic flatworm *S. mansoni* (Weeks et al. 2018) and induction of seizures, c-shapes, and screw-like hyperkinesia at low concentrations (1 µM-10 µM) in the freshwater planarian *Dugesia tigrina* (Thumé and Frizzo 2017), with 10 µM causing lesions and death in *D. tigrina* at 72 hours. We also observed death at 10 µM in both adult and regenerating planarians but did not observe any significant changes in morphology. However, it is possible that the observed body shape phenotypes are short-acting and thus would be missed by only screening the worms at days 7 and 12.

Increases in motility (hyperactivity) have been reported after exposure to fluoxetine (Patocka and Ribeiro 2013; Zewde et al. 2018; Ofoegbu et al. 2019b, a; Duguet et al. 2020) and forskolin (Matsuyama et al. 2004; de Saram et al. 2013; Hirst et al. 2016) in free-living and parasitic flatworms. We also observed hyperactivity with these chemicals in our HTS chronic exposure paradigm here. Most of the chronic effects of fluoxetine observed in this study were in regenerating planarians, whereas past studies have primarily used adult organisms. However, one study has found decreased transformation of somules of the parasitic flatworm *S. mansoni* from the miracidium stage to the primary sporocyst stage after exposure to 2 µM fluoxetine (Taft et al. 2010). *S. mansoni* (Patocka and Ribeiro 2013; Duguet et al. 2020) and *S. mediterranea* (Ofoegbu et al. 2019b, a) exhibited increased motility in response to exposure to fluoxetine across varying time-scales. We also found increased speed and decreased resting at 0.316 µM fluoxetine in regenerating worms.

Exposure to forskolin has been found to decrease transformation from the miracidium stage to the primary sporocyst stage in *S. mansoni* (Kawamoto et al. 1989; Taft et al. 2010). Similarly, we found that chronic exposure to 100 µM forskolin induced lesions at day 7 in regenerating planarians which preceded death at day 12. As observed previously (Ireland et al. 2022), lesions appear to be an early indicator of systemic toxicity. Histamine (180 µM) has been found to induce morphological distortions (primarily supernumerary eyes) in regenerating trunk and tail fragments of *Dugesia lugubris* (Csaba and Bierbauer 1974). We did not observe any effects on eye regeneration at up to 100 µM histamine, though we did find an increase in c-shapes and contraction on day 12 in 100 µM histamine-exposed regenerating *D. japonica*.

In summary, we find that our morphological and behavioral planarian HTS platform allows us to detect previously reported phenotypic changes induced by these compounds with robust, quantitative readouts, in a fraction of the time, and with the unique ability to distinguish between effects on the adult and the developing brain in a single experiment. Moreover, for adult planarians, additional screening could be performed to evaluate acute effects (except for noxious heat sensation, which has negative health consequences and thus is only evaluated on the last screening day); regenerating worms are largely immobile initially, hampering behavioral assays, but morphological changes could still be evaluated.

### 4.3 Efficacy versus toxicity

Parsing out efficacy from toxicity at sublethal concentrations for chronic exposure conditions is difficult, especially since the pharmacology of these compounds in planarians is not understood. Chronic exposure scenarios are important to study, however, because these chemicals are being used for weeks, months, or years. Naively, we expected to see phenotypic patterns reflecting that compounds with a shared mechanism of action will activate (agonists) or block (antagonist) similar molecular pathways. However, except for the few instances highlighted in the Results (Section 3), there was not a clear pattern within effects caused by agonistic and antagonistic drug pairs. Partially, this may be a question of concentration; since we expect effects to be concentration-dependent, it may be difficult to ascertain the observed antagonistic profiles when summarizing effects across concentrations, as done here. For example, mianserin showed several hyperactive hits at low concentrations in regenerating planarians, but only hypoactive hits at higher concentrations. Mianserin has a complex pharmacology as it can act as an antagonist at histamine receptors, serotonin (5-HT) receptors and α2 adrenoceptors, with varying levels of affinity (Pinder 2009) and has been shown to reduce the activity of 5-HT receptors in *D. japonica* (Chan et al. 2016). Thus, depending on the concentration, mianserin could be differentially targeting these various receptors, leading to changes in phenotypic outcomes. It is possible that these hyperactive hits for mianserin, as well as the ones observed with forskolin and fluoxetine, are indicative of desired neuroactive effects whereas the hypoactive effects reflect neurotoxicity. Timing can also be an important factor here, as we would expect neuroactive effects to manifest quicker than adverse effects; however, both are determined by compound metabolism, which is unknown in planarians.

One limitation of our HTS paradigm is static chemical exposure, which does not accurately capture human consumption of these chemicals, which would be dynamic, with regular, repeated dosing. If chemicals were replaced daily, the sensitivity would likely be increased; however, it is unclear whether the endpoints affected would change or the responses would simply be shifted to lower concentrations. Having a direct comparison of these two exposure scenarios would be an interesting future avenue for specific case studies but daily solution changes are clearly not practical for HTS of many chemicals, whose purpose is to provide rapid and efficient hazard identification and prioritization of compounds for further in-depth studies.

## Acknowledgements

The authors would like to thank Veronica Bochenek for help with data analysis. Kevin Bayingana was funded by the Tarble Summer Research Fellowship. Elizabeth Rosenthal was funded by the Frances Velay Women’s Science Summer Research Fellowship from the Panaphil Foundation.

## References

American College of Obstetricians (2010) ACOG CommitteeOpinion No. 462: Moderate caffeine consumption during pregnancy. Obstet Gynecol 116:467–468. https://doi.org/10.1097/AOG.0B013E3181EEB2A1

Bal-Price AK, Hogberg HT, Buzanska L, Coecke S (2010) Relevance of in vitro neurotoxicity testing for regulatory requirements: challenges to be considered. Neurotoxicol Teratol 32:36–41. https://doi.org/10.1016/J.NTT.2008.12.003

Best JB, Morita M (1982) Planarians as a model system for in vitro teratogenesis studies. Teratog Carcinog Mutagen 2:277–291. https://doi.org/10.1002/1520-6866(1990)2:3/4<277::AID-TCM1770020309>3.0.CO;2-8

Bigbee JW, Sharma K V, Chan EL, Bögler O (2000) Evidence for the direct role of acetylcholinesterase in neurite outgrowth in primary dorsal root ganglion neurons. Brain Res 861:354–362. https://doi.org/10.1016/S0006-8993(00)02046-1

Billington CK, Penn RB, Hall IP (2017) β2-agonists. Handb Exp Pharmacol 237:23. https://doi.org/10.1007/164_2016_64

Chan JD, Agbedanu PN, Zamanian M, et al (2014) ‘Death and Axes’: Unexpected Ca2+ Entry Phenologs Predict New Anti-schistosomal Agents. PLOS Pathog 10:e1003942. https://doi.org/10.1371/JOURNAL.PPAT.1003942

Chan JD, Grab T, Marchant JS (2016) Kinetic profiling an abundantly expressed planarian serotonergic GPCR identifies bromocriptine as a perdurant antagonist. Int J Parasitol Drugs Drug Resist 6:356–363. https://doi.org/10.1016/j.ijpddr.2016.06.002

Chang EA, Beyhan Z, Yoo MS, et al (2009) Increased cellular turnover in response to fluoxetine in neuronal precursors derived from human embryonic stem cells. Int J Dev Biol 54:707–715. https://doi.org/10.1387/IJDB.092851EC

Chen L, Bell EM, Browne ML, et al (2014) Exploring maternal patterns of dietary caffeine consumption before conception and during pregnancy. Matern Child Health J 18:2446–2455. https://doi.org/10.1007/S10995-014-1483-2

Chifiriuc MC, Ratiu AC, Popa M, Ecovoiu A Al (2016) Drosophotoxicology: An Emerging Research Area for Assessing Nanoparticles Interaction with Living Organisms. Int J Mol Sci 17:. https://doi.org/10.3390/IJMS17020036

Cipriani A, Furukawa TA, Salanti G, et al (2009) Comparative efficacy and acceptability of 12 new-generation antidepressants: a multiple-treatments meta-analysis. Lancet 373:746–758. https://doi.org/10.1016/S0140-6736(09)60046-5

Cochet-Escartin O, Mickolajczk KJ, Collins E-MS (2015) Scrunching: a novel escape gait in planarians. Phys Biol 12:055001. https://doi.org/doi:10.1088/1478-3975/12/5/056010

Csaba G, Bierbauer J (1974) Investigations on the specificity of hormone receptors in planarians. Gen Comp Endocrinol 22:132–134. https://doi.org/10.1016/0016-6480(74)90095-1

Currie KW, Pearson BJ (2013) Transcription factors lhx1/5-1 and pitx are required for the maintenance and regeneration of serotonergic neurons in planarians. Dev 140:3577–3588. https://doi.org/10.1242/dev.098590

Daw JR, Hanley GE, Greyson DL, Morgan SG (2011) Prescription drug use during pregnancy in developed countries: a systematic review. Pharmacoepidemiol Drug Saf 20:895–902. https://doi.org/10.1002/PDS.2184

de Saram PSR, Ressurreição M, Davies AJ, et al (2013) Functional Mapping of Protein Kinase A Reveals Its Importance in Adult Schistosoma mansoni Motor Activity. PLoS Negl Trop Dis 7:. https://doi.org/10.1371/journal.pntd.0001988

Dubovicky M, Belovicova K, Csatlosova K, Bogi E (2017) Risks of using SSRI / SNRI antidepressants during pregnancy and lactation. Interdiscip Toxicol 10:30. https://doi.org/10.1515/INTOX-2017-0004

Duguet TB, Glebov A, Hussain A, et al (2020) Identification of annotated bioactive molecules that impair motility of the blood fluke Schistosoma mansoni. Int J Parasitol Drugs Drug Resist 13:73–88. https://doi.org/10.1016/j.ijpddr.2020.05.002

Ellman GL, Courtney KD, Andres V, Featherstone RM (1961) A new and rapid colorimetric determination of acetylcholinesterase activity. Biochem Pharmacol 7:88–95. https://doi.org/10.1016/0006-2952(61)90145-9

Ferrero S, Colombo BM, Ragni N (2004) Maternal arrhythmias during pregnancy. Arch Gynecol Obstet 269:244–253. https://doi.org/10.1007/S00404-002-0461-X

Giacomotto J, Ségalat L (2010) High-throughput screening and small animal models, where are we? Br J Pharmacol 160:204. https://doi.org/10.1111/J.1476-5381.2010.00725.X

Haas DM, Marsh DJ, Dang DT, et al (2018) Prescription and Other Medication Use in Pregnancy. Obstet Gynecol 131:789. https://doi.org/10.1097/AOG.0000000000002579

Hagstrom D, Cochet-Escartin O, Collins E-MS (2016) Planarian brain regeneration as a model system for developmental neurotoxicology. Regeneration 3:65–77. https://doi.org/10.1002/reg2.52

Hagstrom D, Cochet-Escartin O, Zhang S, et al (2015) Freshwater planarians as an alternative animal model for neurotoxicology. Toxicol Sci 147:270–285. https://doi.org/10.1093/toxsci/kfv129

Hagstrom D, Hirokawa H, Zhang L, et al (2017) Planarian cholinesterase: in vitro characterization of an evolutionarily ancient enzyme to study organophosphorus pesticide toxicity and reactivation. Arch Toxicol 91:2837–2847. https://doi.org/10.1007/s00204-016-1908-3

Hagstrom D, Zhang S, Ho A, et al (2018a) Planarian cholinesterase: molecular and functional characterization of an evolutionarily ancient enzyme to study organophosphorus pesticide toxicity. Arch Toxicol 92:1161–1176. https://doi.org/10.1007/s00204-017-2130-7

Hagstrom D, Zhang S, Ho A, et al (2018b) Planarian cholinesterase: molecular and functional characterization of an evolutionarily ancient enzyme to study organophosphorus pesticide toxicity. Arch Toxicol 92:1161–1176. https://doi.org/10.1007/s00204-017-2130-7

Hansen C, Desrosiers TA, Wisniewski K, et al (2020) Use of antihistamine medications during early pregnancy and selected birth defects: The National Birth Defects Prevention Study, 1997–2011. Birth Defects Res 112:1234–1252. https://doi.org/10.1002/BDR2.1749

Helmcke KJ, Avila DS, Aschner M (2010) Utility of Caenorhabditis elegans in high throughput neurotoxicological research. Neurotoxicol Teratol 32:62–67. https://doi.org/10.1016/J.NTT.2008.11.005

Hirst NL, Lawton SP, Walker AJ (2016) Protein kinase A signalling in Schistosoma mansoni cercariae and schistosomules. Int J Parasitol 46:425–437. https://doi.org/10.1016/j.ijpara.2015.12.001

Hunt PR, Camacho JA, Sprando RL (2020) Caenorhabditis elegans for predictive toxicology. Curr Opin Toxicol 23–24:23–28. https://doi.org/10.1016/J.COTOX.2020.02.004

Ireland D, Bochenek V, Chaiken D, et al (2020) Dugesia japonica is the best suited of three planarian species for high-throughput toxicology screening. Chemosphere 253:126718. https://doi.org/10.1016/j.chemosphere.2020.126718

Ireland D, Zhang S, Bochenek V, et al (2022) Differences in neurotoxic outcomes of organophosphorus pesticides revealed via multi-dimensional screening in adult and regenerating planarians. bioRxiv 496617

Joglar JA, Page RL (2012) Treatment of Cardiac Arrhythmias During Pregnancy. Drug Saf 1999 201 20:85–94. https://doi.org/10.2165/00002018-199920010-00008

Kawamoto F, Shozawa A, Kumada N, Kojima K (1989) Possible roles of cAMP and Ca2+ in the regulation of miracidial transformation in Schistosoma mansoni. Parasitol Res 75:368–374. https://doi.org/10.1007/BF00931132

Kokel D, Rennekamp AJ, Shah AH, et al (2012) Behavioral barcoding in the cloud: embracing data-intensive digital phenotyping in neuropharmacology. Trends Biotechnol 30:421–425. https://doi.org/10.1016/j.tibtech.2012.05.001

Layer PG, Klaczinski J, Salfelder A, et al (2013) Cholinesterases in development: AChE as a firewall to inhibit cell proliferation and support differentiation. Chem Biol Interact 203:269–276. https://doi.org/10.1016/j.cbi.2012.09.014

Li MH (2013) Acute toxicity of 30 pharmaceutically active compounds to freshwater planarians, Dugesia japonica. Toxicol Environ Chem 95:1157–1170. https://doi.org/10.1080/02772248.2013.857671

Matsuyama H, Takahashi H, Watanabe K, et al (2004) The involvement of cyclic adenosine monophosphate in the control of Schistosome miracidium cilia. J Parasitol 90:8–14. https://doi.org/10.1645/GE-52R1

Moustakas D, Mezzio M, Rodriguez BR, et al (2015) Guarana provides additional stimulation over caffeine alone in the planarian model. PLoS One 10:. https://doi.org/10.1371/journal.pone.0123310

Müller TC, Rocha JBT, Morsch VM, et al (2002) Antidepressants inhibit human acetylcholinesterase and butyrylcholinesterase activity. Biochim Biophys Acta 1587:92–98. https://doi.org/10.1016/S0925-4439(02)00071-6

OECD (2021) Guideline No. 497: Defined Approaches on Skin Sensitisation. In: OECD Guidelines for the Testing of Chemicals. OECD Publishing, Paris

Ofoegbu PU, Campos D, Soares AMVM, Pestana JLT (2019a) Combined effects of NaCl and fluoxetine on the freshwater planarian, Schmidtea mediterranea (Platyhelminthes: Dugesiidae). Environ Sci Pollut Res 26:11326–11335. https://doi.org/10.1007/s11356-019-04532-4

Ofoegbu PU, Lourenço J, Mendo O, et al (2019b) Effects of low concentrations of psychiatric drugs (carbamazepine and fluoxetine) on the freshwater planarian, Schmidtea mediterranea. Chemosphere 217:542–549. https://doi.org/10.1016/j.chemosphere.2018.10.198

Paraoanu LE, Steinert G, Klaczinski J, et al (2006) On functions of cholinesterases during embryonic development. J Mol Neurosci 30:201–4. https://doi.org/10.1385/JMN:30:1:201

Patocka N, Ribeiro P (2013) The functional role of a serotonin transporter in Schistosoma mansoni elucidated through immunolocalization and RNA interference (RNAi). Mol Biochem Parasitol 187:32–42. https://doi.org/10.1016/j.molbiopara.2012.11.008

Peterson RT, Nass R, Boyd WA, et al (2008) USE OF NON-MAMMALIAN ALTERNATIVE MODELS FOR NEUROTOXICOLOGICAL STUDY. Neurotoxicology 29:546. https://doi.org/10.1016/J.NEURO.2008.04.006

Pinder RM (2009) Mianserin: Pharmacological and clinical correlates. http://dx.doi.org/103109/08039489109096678 45:13–26. https://doi.org/10.3109/08039489109096678

Rand MD (2010) Drosophotoxicology: the growing potential for Drosophila in neurotoxicology. Neurotoxicol Teratol 32:74. https://doi.org/10.1016/J.NTT.2009.06.004

Rawls SM, Patil T, Yuvasheva E, Raffa RB (2010) First evidence that drugs of abuse produce behavioral sensitization and cross sensitization in planarians. Behav Pharmacol 21:301–313. https://doi.org/10.1097/FBP.0b013e32833b0098

Ruszkiewicz JA, Pinkas A, Miah MR, et al (2018) C. elegans as a model in developmental neurotoxicology. Toxicol Appl Pharmacol 354:126–135. https://doi.org/10.1016/J.TAAP.2018.03.016

Sabry Z, Ho A, Ireland D, et al (2019) Pharmacological or genetic targeting of Transient Receptor Potential (TRP) channels can disrupt the planarian escape response. PLoS One 14:e0226104. https://doi.org/10.1371/journal.pone.0226104

Ségalat L (2007) Invertebrate animal models of diseases as screening tools in drug discovery. ACS Chem Biol 2:231–236. https://doi.org/10.1021/CB700009M

Servey J, Chang J (2014) Over-the-Counter Medications in Pregnancy - American Family Physician. Am Fam Physician 90:548–555

Shettigar N, Chakravarthy A, Umashankar S, et al (2021) Discovery of a body-wide photosensory array that matures in an adult-like animal and mediates eye-brain-independent movement and arousal. Proc Natl Acad Sci U S A 118:. https://doi.org/10.1073/PNAS.2021426118

So M, Bozzo P, Inoue M, Einarson A (2010) Safety of antihistamines during pregnancy and lactation. Can Fam Physician 56:427

Taft AS, Norante FA, Yoshino TP (2010) The identification of inhibitors of Schistosoma mansoni miracidial transformation by incorporating a medium-throughput small-molecule screen. Exp Parasitol 125:84–94. https://doi.org/10.1016/j.exppara.2009.12.021

Talbot JA, Currie KW, Pearson BJ, Collins EMS (2014) Smed-dynA-1 is a planarian nervous system specific dynamin 1 homolog required for normal locomotion. Biol Open 3:627–634. https://doi.org/10.1242/bio.20147583

The Canadian Preterm Labor Investigators Group (1992) Treatment of Preterm Labor with the Beta-Adrenergic Agonist Ritodrine. N Engl J Med 327:308–312. https://doi.org/10.1056/NEJM199207303270503

Thumé IS, Frizzo ME (2017) Sertraline Induces Toxicity and Behavioral Alterations in Planarians. Biomed Res Int 2017:. https://doi.org/10.1155/2017/5792621

Weeks JC, Roberts WM, Leasure C, et al (2018) Sertraline, Paroxetine, and Chlorpromazine Are Rapidly Acting Anthelmintic Drugs Capable of Clinical Repurposing. Sci Rep 8:. https://doi.org/10.1038/s41598-017-18457-w

Witter FR, Zimmerman AW, Reichmann JP, Connors SL (2009) In utero beta 2 adrenergic agonist exposure and adverse neurophysiologic and behavioral outcomes. Am J Obstet Gynecol 201:553–559. https://doi.org/10.1016/J.AJOG.2009.07.010

Wu JP, Li MH (2018) The use of freshwater planarians in environmental toxicology studies: Advantages and potential. Ecotoxicol Environ Saf 161:45–56. https://doi.org/10.1016/j.ecoenv.2018.05.057

Yang M, Liu S, Hu L, et al (2018) Effects of the antidepressant, mianserin, on early development of fish embryos at low environmentally relevant concentrations. Ecotoxicol Environ Saf 150:144–151. https://doi.org/10.1016/J.ECOENV.2017.12.024

Zewde AM, Yu F, Nayak S, et al (2018) PLDT (planarian light/dark test): an invertebrate assay to quantify defensive responding and study anxiety-like effects. J Neurosci Methods 293:284–288. https://doi.org/10.1016/j.jneumeth.2017.10.010

Zhang S, Hagstrom D, Hayes P, et al (2019a) Multi-behavioral endpoint testing of an 87-chemical compound library in freshwater planarians. Toxicol Sci 167:105–115. https://doi.org/10.1093/toxsci/kfy145

Zhang S, Ireland D, Sipes NS, et al (2019b) Screening for neurotoxic potential of 15 flame retardants using freshwater planarians. Neurotoxicol Teratol 73:54–66. https://doi.org/10.1016/j.ntt.2019.03.003

