## Supplemental Figure 1 for "Adult and regenerating planarians respond differentially to chronic drug exposure"

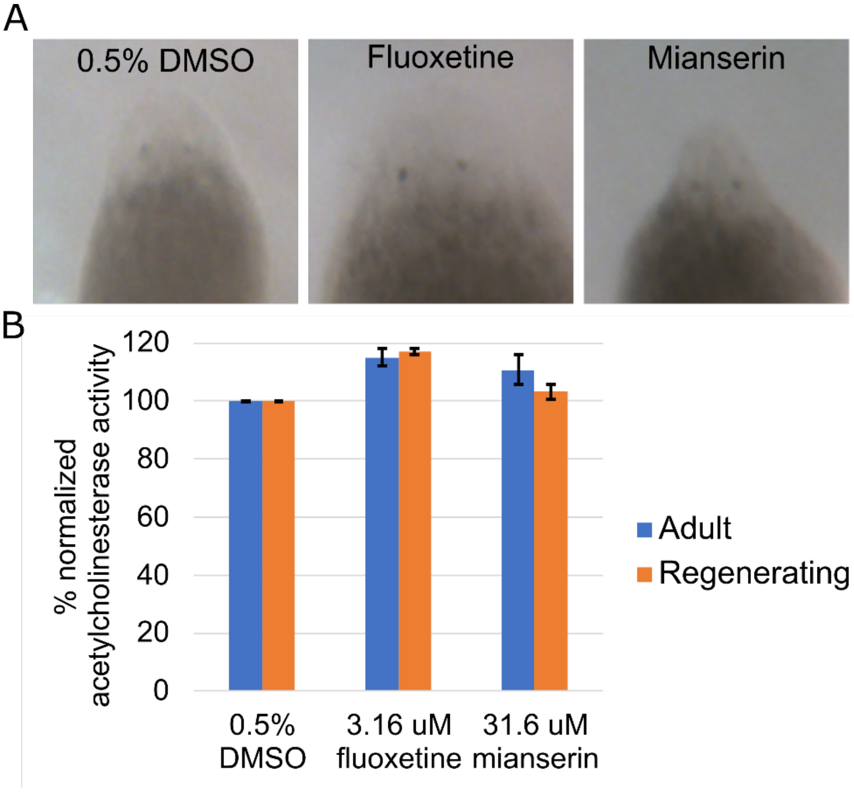


**Supplementary Figure 1.** Fluoxetine and mianserin do not affect eye regeneration or acetylcholinesterase activity. A) Eyes were visible by day 5 in regenerating planarians exposed to 0.5% DMSO, 3.16 µM fluoxetine or 3.16 µM mianserin. Note that 3.16 µM mianserin can induce contraction in regenerating planarians (Figure 5A). Representative images of the highest tested concentrations are shown. n=10. B) Ellman assays were used to measure acetylcholinesterase activity in adult and regenerating planarians exposed for 12 days. Error bars indicate standard error of 6 technical replicates from 2 independent experiments (biological replicates).
