## Supplemental Table 1 for "Adult and regenerating planarians respond differentially to chronic drug exposure"

**Supplemental Table**: Summary of past findings in the literature of the effects of the drugs tested here on various planarian species and *Schistosoma mansoni*. N/A: information not available. Concentrations were converted to µM to aid comparison.

| Chemical | Reference | Species | LOEL | Exposure Duration[e] | Endpoints Affected |
|---|---|---|---|---|---|
| Forskolin | (Hirst, Lawton, and Walker 2016) | *Schistosoma mansoni* (somules) | 50 µM[a] | Acute | contractile hyperkinesia |
| | (de Saram et al. 2013) | *S. mansoni* | 50 µM[b] | Acute | ↑ contractions (gross random muscular movements), coiling, body bends, thrashing (overall – hyperkinetic) |
| | (Taft, Norante, and Yoshino 2010) | *S. mansoni* | 10-20 µM[a] | Short-term | ↓ transformation from the miracidium stage to the primary sporocyst stage |
| | (Matsuyama et al. 2004) | *S. mansoni* | 0.5 µM | Acute | ↑ motility (swimming) (only in hypertonic NaCl solution) |
| | (Kawamoto et al. 1989) | *S. mansoni* | 50 µM | Acute | ↓ transformation from the miracidium stage to the primary sporocyst stage |
| MDL-12,330A | (Matsuyama et al. 2004) | *S. mansoni* | 50 µM[b] | Acute | ↓ motility (swimming) |
| Caffeine | (Moustakas et al. 2015) | *Dugesia tigrina* | 10 mM | Acute | ↓ motility (pLMV), ↑ contractions |
| | (M. H. Li 2013) | *D. japonica* | 5.1 mM (24 h), 3.3 mM (48 h), 2.5 mM (72 h), 1.9 mM (96h) [c] | Subacute, Short-term, chronic | ↑ lethality |
| | (Rawls et al. 2010) | *Dugesia dorotocephala* | 1 mM | Acute | ↑ C-like hyperkinesia |
| | | *D. dorotocephala* | 257 µM | Not specified | ↑ locomotor activity |

| | | | | | |
|---|---|---|---|---|---|
| | (Best and Morita 1982) | | 515 µM | Not specified | ↓ locomotor activity, ↑ head lesions, resorptions |
| Histamine | (Csaba and Bierbauer 1974) | *Dugesia lugubris* (regenerating fragments) | 180 µM[a] | Chronic | ↑ morphological distortions (supernumerary eyes) |
| Mianserin | (Shettigar et al. 2021) | *S. mediterranea* (regenerating tail fragments) | 60 µM[a] | Acute | ↑ peristaltic, muscle-based movement in response to UV-A stimulation |
| | (Chan, Grab, and Marchant 2016) | *D. japonica* | 10 µM[a] | Acute | ↓ mobility |
| | (Chan et al. 2014) | *D. japonica* (regenerating trunk fragments) | 10 µM[a] | Short-term | ↑ two-headed planarians |
| | | *S. mansoni* (somules) | 5 µM[b] | Acute | ↓ contractility |
| | (Currie and Pearson 2013) | *S. mediterranea* | 20 µM[a] | Acute | ↓ gliding, ↑ inchworm-like crawling, ↑ uncoordinated motile cilia, ↓ ciliary beat frequency |
| Fluoxetine | (Duguet et al. 2020) | *S. mansoni* | 1 µM[b] | Subacute, chronic | ↑ motility |
| | | | 10 µM[a] | Chronic | ↑ elongated phenotype |
| | (Taft, Norante, and Yoshino 2010) | *S. mansoni* | 20 µM[a] | Short-term | ↓ transformation from the miracidium stage to the primary sporocyst stage |
| | (Patocka and Ribeiro 2013) | *S. mansoni* | 100 µM[a] | Acute | ↑ motility |
| | (Ofoegbu, Campos, et al. 2019) | *S. mediterranea* | 30 nM | Chronic | ↑ locomotion (pLMV), ↓ fissioning |
| | | *S. mediterranea* | 0.5 µM[c] | Chronic | ↑ lethality |

| | | | | | |
|---|---|---|---|---|---|
| | Ofoegbu, Lourenço, et al. 2019) | | 3 nM | Chronic | ↑ locomotion, ↓ feeding |
| | | | 30 nM | Chronic | ↓ fissioning |
| | | | 300 pM | Chronic | ↑ DNA damage |
| | (Zewde et al. 2018) | *D. dorotocephala* | 1 µM | Acute | ↑ time spent in the light |
| Sertraline | (Weeks et al. 2018) | *S. mansoni* | 5 µM | Subacute, Short-term | ↑ rounding and degeneration, ↓ motility |
| | | | 8.43 µM[d] | Subacute | ↓ motility |
| | (Thumé and Frizzo 2017) | *D. tigrina* | 0.4 µM | Acute | ↓ locomotion (pLMV) |
| | | | 1 µM | Subacute | C-shape, screw-like hyperkinesia |
| | | | 4 µM | Acute, Subacute | ↑ seizures , ↓ locomotion (pLMV), |
| | | | 10 µM | Short-term | Severe lesions and death |

[a]Only concentration tested
[b]Lowest concentration tested
[c]$LC_{50}$
[d]$EC_{50}$
[e]Exposure durations were separated into three groups: Acute (≤3 hours), Subacute (3-24 hours), Short-term (2-3 days), and Chronic (>3 days)
